# SatXplor – A comprehensive pipeline for satellite DNA analyses in complex genome assemblies

**DOI:** 10.1101/2024.08.09.607335

**Authors:** Marin Volarić, Nevenka Meštrović, Evelin Despot-Slade

**Affiliations:** Ruđer Bošković Institute, Zagreb, Croatia

## Abstract

Satellite DNAs (satDNAs) are tandemly repeated sequences that make up a significant portion of almost all eukaryotic genomes. Although satDNAs have been shown to play a very important role in genome organization and evolution, they are relatively poorly analysed even in model. One of the main reasons for the current lack of in-depth studies on satDNAs is their underrepresentation in genome assemblies. The complexity and highly repetitive nature of satDNAs make their analysis challenging, and there is a need for efficient tools that can ensure accurate annotation and analysis of satDNAs.

We present a novel pipeline, named Satellite DNA Exploration (SatXplor), designed to robustly characterize satDNA elements and analyse their arrays and flanking regions. SatXplor is benchmarked against curated satDNA datasets from diverse species, showcasing its versatility across genomes with varying complexities and different satDNA profile. Component algorithms excel in the identification of tandemly repeated sequences and for the first time enable evaluation of satDNA variation and array annotation with the addition of information about surrounding genomic landscape.

SatXplor is an innovative pipeline for satDNA analysis that can be paired with any tool used for satDNA detection, offering insights into the structural characteristics, array determination and genomic context of satDNA elements. By integrating various computational techniques, from sequence analysis and homology investigation to advanced clustering and graph-based methods, it provides a versatile and comprehensive approach to explore the complexity of satDNA organization and to understand the underlying mechanisms and evolutionary aspects. It is open-source and freely accessible at https://github.com/mvolar/SatXplor.

## Introduction

Repetitive DNA sequences comprise a substantial portion of the eukaryotic genomes and play pivotal roles in genome stability, evolution and functional diversification (Biscotti et al., 2015). Based on their abundance, they can be divided into two main classes: tandemly repeated satellite DNAs (satDNAs) and dispersed repetitive DNAs such as transposable elements (TEs) (reviewed in Liao et al., 2023). SatDNA constitutes a unique class of repetitive elements characterized by tandemly repeated units, also called monomers. They are generally characterized by monomer lengths of 150–400 bp (reviewed in Garrido-Ramos, 2017) and long arrays which can be up to 1 Mb in length. SatDNAs are an integral part of the (peri)centromeric region of almost all eukaryotes but recently have also been increasingly found in euchromatin (Cabral-de-Mello et al., 2023; Pavlek et al., 2015; Rico-Porras et al., 2024). They are among the fastest evolving parts of the eukaryotic genome, leading to species-specific satDNA profiles. Moreover, in certain species, the presence of more than hundreds of distinct satDNA families underscores their remarkable diversity (Utsunomia et al., 2019). Also, satDNA can form higher-order repeat (HOR) organization through specific arrangement of different satDNA monomers creating structured repeated pattern further adding complexity to these regions (Sujiwattanarat et al., 2015). Due to the variability of the monomers, possibly very long satDNA arrays, high genome abundance and the presence of many different satDNA families in the genome, satDNAs cause problems in genome assembly based on the most commonly used Illumina sequencing. Consequently, they are not or only insufficiently represented in genome assemblies.

A new technology development of Pacific Biosciences (PacBio) and Oxford Nanopore Technologies (ONT) long-read sequencing has made it possible to capture long satDNA regions for the first time. Existing genome assembly algorithms based on one or more main principles such as overlap-layout-consensus (OLC) and graph-based computation often produce contiguous assemblies in genomic regions. However, to specifically target obstacles that arise with high genome repetitiveness, hybrid approaches are often needed to further improve and increase the repeat resolution using ultra long reads (Wee et al., 2019). Obtained assemblies are becoming more complete, even reaching telomere to telomere level recently achieved for human genome including (peri)centromere (Nurk et al., 2022). Thus, approaches coupling long read sequencing with assembly algorithms which allow repetitive DNA enrichment, provide a great platform for in-depth assessment of satDNAs in any species of interest.

Today, many different tools have been developed specifically for detection and identification of repetitive DNA in sequenced genomic data. For example, the assembly-free algorithm TAREAN performs graph-based clustering for high throughput detection of satDNAs on short Illumina reads (Novák et al., 2017). The data obtained from this analysis contains only basic information about satDNA, such as the variability and abundance of monomers but without any information on genome organization. The other frequently used program, Tandem Repeat Finder (TRF) offers detection of tandem repeats on assembled sequences by recognizing periodicity and providing monomers and their consensus (Benson, 1999). Mreps utilizes the Hamming distance approach for satDNA detection on whole genomes (Kolpakov, 2003), but it has limitations when investigating complex genomic regions. There is also a RepBase database (Bao et al., 2015) of several classes of repetitive elements, including satDNA with curated annotations and classifications. It is used by RepeatMasker (Smit, AFA, Hubley, R & Green, 2015) to accurately identify previously described repeats relying on sequence similarity and evolutionary conservation. Recently there have been developed several tools such as TRASH (Wlodzimierz et al., 2023) and NanoTRF (Kirov et al., 2022) that provide improved detection and annotation of satDNA in long-read genome assemblies. However, what all these programs for satDNA detection lack is the capability of finding a complete set of monomers on the genome scale, accurate detection of arrays and surrounding regions as well as revealing their dynamics. Therefore, proper genome-wide annotation of satDNA monomers and arrays is a prerequisite for disclosing the organization, mechanisms of propagation, and evolutionary processes for sequences in question.

Going past the detection step, any comprehensive analysis requires extensive understanding of both bioinformatics and repetitive DNA behaviour. This is especially true for satDNAs which are often investigated only at the monomer level with little insight into their array organization and genomic environment. The absence of a unified methodology for satDNA analyses on the genome scale leads to diverse approaches across literature often focusing only on specific organisms, satDNA families or even one satDNA sequence (Ruiz-Ruano et al., 2016; Sproul et al., 2020; Vondrak et al., 2020).

The flour beetle *Tribolium castaneum* as a model genome for satDNA analyses offers unique advantage as there are well-defined satDNA families present in euchromatin (Pavlek et al., 2015) in a highly repetitive genome with one satDNA occupying large centromeres (Gržan et al., 2020) together with a well described satellitome (Gržan et al., 2023). Our recent study disclosing mechanisms of euchromatic satDNA spread in *T. castaneum* is based on an improved genome assembly using nanopore sequencing and its enrichment in the repetitive part of satDNA in particular (Volarić et al., 2024). Investigation of euchromatic satDNAs behaviour proved to be challenging due to their specific characteristics, different genome proportions, varying array lengths (1-100 kb), and distribution across multiple genomic loci. Solving these complexities in the study required the development of customized scripts for the annotation and analysis of both satDNA monomers and arrays, the precise definition of array ends and the analysis of their surrounding regions.

In this work, we improved and automated the approach initially used in *T. castaneum* analyses in an integrated pipeline – SatXplor. It encompasses multiple algorithms for thorough satDNA exploration and analyses validated on different genomes utilizing multiple publicly available long read based assemblies with well-described satDNAs. To validate and further develop the SatXplor pipeline, we selected moderately repetitive and smaller genomes (150-350 Mb) of *Meloidogyne* nematodes (*M. incognita, M. arenaria*) and *Drosophila melanogaster*, highly repetitive and very large genome (6.3 Gb) of the locust *Locusta migratoria*, while the genome of *Arabidopsis thaliana* served as a plant model. SatXplor represents a unique pipeline of powerful tools for in-depth exploration of middle and lower copy satDNAs, providing a standardized and comprehensive solution for satDNA analysis that enables deeper insights into genome organization and the potential mechanisms underlying these regions in any long-read based genome assembly.

## Results

The SatXplor pipeline contains several algorithms for satDNA exploration and analysis (Figure 1) requiring only FASTA sequences of potential satDNAs and a long read based assembly. The first step is the accurate and complete detection of satDNA monomers in respective assemblies using a BLAST search followed by a homology-based inspection. The next algorithm recognizes satDNA organization followed by array definition and precise profiling to disclose potential patterns in arrays‘ organization. Next, individual satDNA monomers, arrays and their flanking regions are extracted and serve as input for subsequent analysis. In satDNA monomer analyses PCA (Principal Component Analysis) and UMAP (Uniform Manifold Approximation and Projection) are used to cluster satDNA monomers to disclose their relationship. Then, an undirected graph is created to comprehensively visualize the relationships between monomers in satDNA arrays. A kmer-based analysis increases the precision in determining precise array boundaries allowing extraction of flanking regions. Arrays boundaries and flanking regions serve as a basis for the detection of potential micro- (using seqLogo) and macrohomologies (using distance maps). Whole output is stored within results folder that is presented in the structure depicted in Suppl. Figure 1.

**Figure 1.**
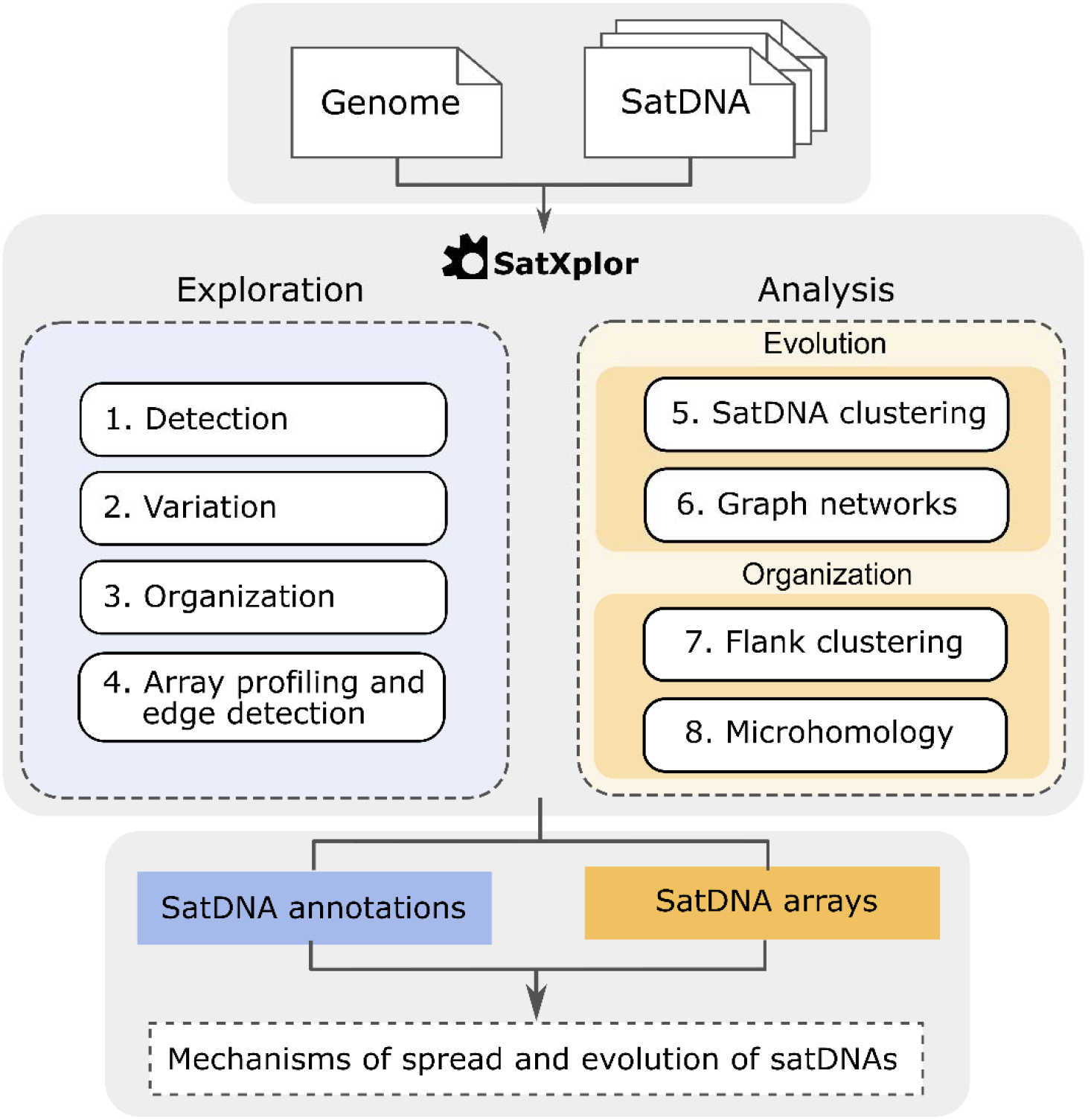
Workflow of SatXplor pipeline for comprehensive satDNA annotation and analysis in long-read-based genome assemblies. SatXplor pipeline consists of four exploration algorithms for finding structural characteristics and variants of satDNA elements together with four analysis algorithms for evolutionary insight and organizational patterns dissecting further the satDNA genomic landscape. The output of SatXplor provides detailed annotations and visualizations, shedding light on the mechanisms underlying their spread and evolution within the genome.

The robustness of SatXplor toolbox for analysis of satDNAs was tested in several species with genomes of various sizes and repetitiveness (Suppl. Table 1) and structurally different satDNAs (Suppl. Table 2). In *D. melanogaster* well described 1.688 satDNA was investigated (de Lima et al., 2020), in *Meloidogyne* we used the database of tandem repeats (Despot-Slade et al., 2021, 2022), in *L. migratoria* we utilized well described satellitome (Ruiz-Ruano et al., 2016) and explored satDNAs of *A. thaliana* described in Simoens et al., 1988. The detection of satDNA consensus monomer sequence for each species was previously performed and experimentally validated using molecular methods and bioinformatics tools (TRF, TAREAN) (Suppl. Table 2).

### 1. SatDNA exploration

#### 1.1. SatDNA detection

First step in SatXplor pipeline is the detection of all potential monomers in the genome, which is performed using NCBI BLAST+ executable and satDNA consensus sequences as query. A phyton-based BLAST wrapper with parameters best suited for satDNA detection (see Materials and methods section) controls the process and generates a result table with all potential hits in the assembly (Suppl. Table 3). This table serves as the basis for subsequent analyses within the satDNA toolbox, enabling the extraction of essential satDNA information. This includes the localization of each satDNA monomer on the target sequence, as well as rough estimates of percentage identity and query coverage. It is noteworthy that, at this stage, no filtering step is applied, allowing for a comprehensive downstream exploration of satDNAs.

#### 1.2. Finding homology between annotated satDNA

The first objective of SatXplor is to visualize the satDNA variability in the assembly. For this purpose, we have developed a visualization technique that quickly and efficiently analyses all detected monomers in the genome. Using 2D relative density approximation on data present in the BLAST output table (Suppl. Table 3), potential groupings of satDNA monomers are visualized, allowing for a clear assessment of intragenomic variability of a satDNA family. This analysis results in different scenarios of monomer conservation of particular satDNA within the genome under investigation (Figure 2A). For instance, satDNA that is characterized by low monomer variability is visible as a single spot in the graph (Figure 2A, Example 1). Slight variation is visible as light areas on the graph (Figure 2A, Example 2), whereas higher variation is depicted as several spots of similar intensity (Figure 2A, Example 3). Finally, some satDNAs may exhibit high variability and degeneracy in the monomer sequence, which becomes visible as a large region of intensive signal that extends toward lower percentages (Figure 2A, Example 4). The information from this graph can also be used to define specific sequence identity and reference sequence coverage parameters to annotate and extract all monomers for particular satDNA from the genome. This comprehensive characterization underscores the utility of this approach and can serve as a basis for any downstream satDNA assessment.

**Figure 2.**
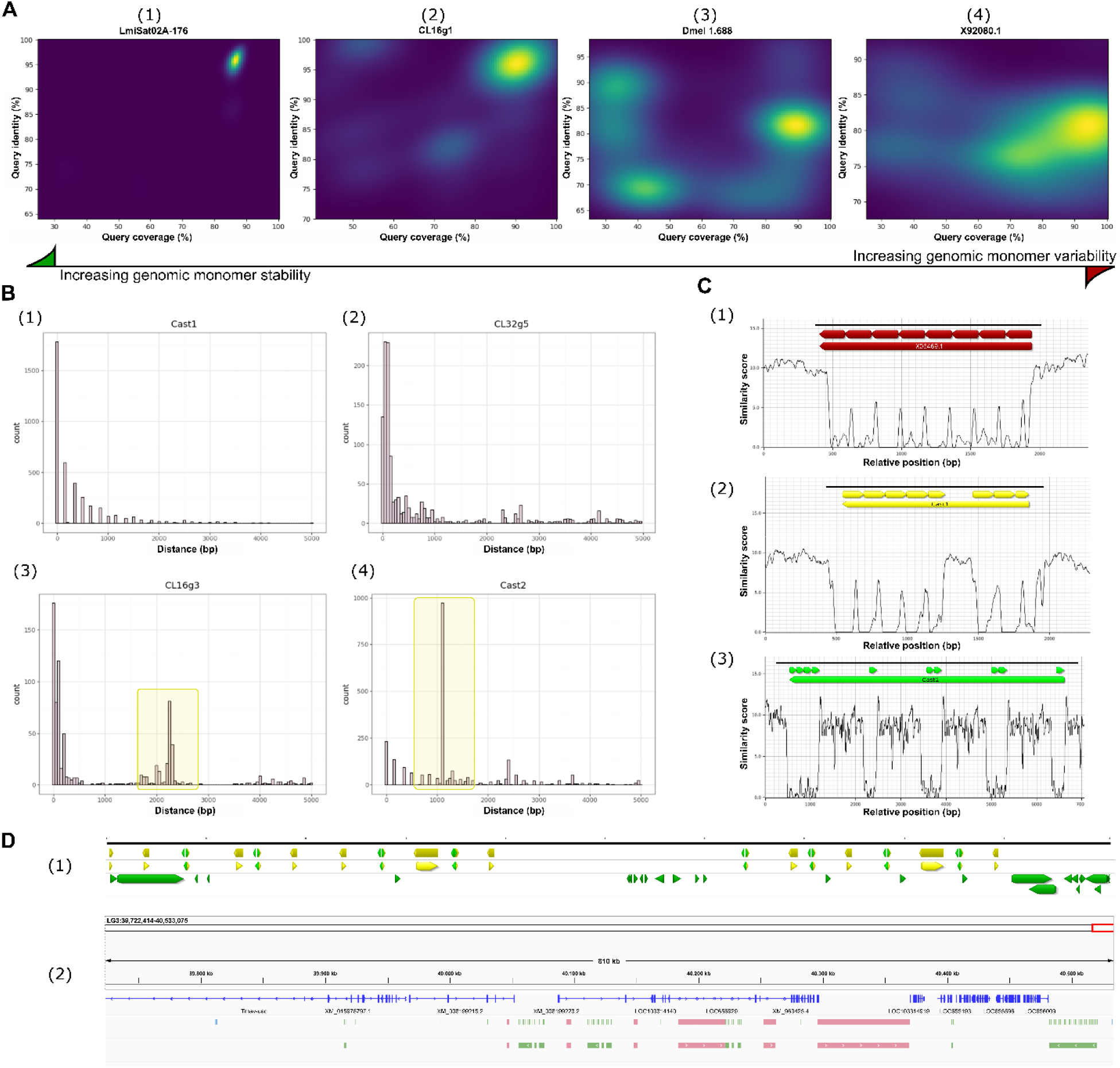
Preparation and annotation of satDNA data with SatXplor. A Representative scenarios of different results for BLAST based monomer detection in genomes of interest. Depending on the satDNA characteristics different levels of monomer variability can be observed from extremely low (1), through moderate (2) and highly variable (3), to extremely variable and fragmented monomers (4). **B SatDNA organization**. Depicted are certain scenarios of tandem organization for specific satDNA monomers. SatDNAs can have low distances between monomers (1,2) with values often being random, indicating homogenous array. This approach can also detect complex organization of satDNA monomers characterized by the interspersion with other specific sequence (highlighted spike in 3, 4). **C Precise array profiling**. Per-array monomer distance profiles generated by the kmer-counting method give powerful visualizations to study patterns of specific arrays ‘ organization. For example, a homogenous tandem array (1), a slightly discontinuous array (2) and a discontinuous array (3) can be observed. Coloured arrows represent satDNA monomers and arrays corresponding to graph profiles beneath. **D Annotation and visualization**. Output of SatXplor can easily be used in different bioinformatics visualization tools such as Geneious (1) and IGV (2). Here, each satDNA monomer and accompanying array are shown in separated tracks and can then later be separated between different satDNA families. All satDNA names and species used (A-D) are listed in Suppl. Table 2.

#### 1.3. Detection of complex satDNA organization

Although we have successfully identified parameters for satDNA detection using the approach described above (Figure 2A), many satDNAs do not form pure tandem arrays, but are rather interspersed with other sequences. Therefore, the crucial step before we start the subsequent determination and analysis of satDNA arrays is the detection of possible interspersed sequences between satDNA monomers. For each satDNA family, SatXplor outputs a plot with the approximate distance to the nearest monomer in a strand-independent manner, finds the peaks and then elongates both the satDNAs and the newly found interspersed sequences into larger arrays, outputting both the annotation and general statistics of the arrays found. For example, satDNA which occurs in homogeneous tandem repeat arrays, the graph shows a dominant single peak at distance values ∼0 (Figure 2B, Example 1 and 2). However, when a sequence is inserted between satDNA monomers represented by the presence of one or more peaks at distances greater than monomer length (Figure 2B, Example 3 and 4). Additional feature of this algorithm is the capability to detect and extract all sequences that come in between satDNA monomers of interest, even if they are not tandemly repeated or if there are multiple types of complex organization for a single satDNA family.

#### 1.4. Kmer-based profiling and edge detection

Defining the ends of satDNA arrays is crucial for downstream analyses of propagation mechanisms and evolutionary dynamics within the genome. The main problem for proper arrays ‘ edge definition is the tendency of arrays to degenerate in sequence at their edges. For example, first or last monomers of an array are often truncated (Sproul et al., 2020) and subject to higher sequence variability. This limits the finding of potential micro- and macrohomologies in flanking regions (Suppl. Figure 2). To solve this problem, we developed a novel method for precise edge detection employing a per-array kmer-distance calculating method followed by postprocessing. The result of this analysis are precisely defined array edges subsequently used in the analysis as well as per-array profiles. Utilizing the kmer-counting method, we generated monomer distance profiles for each detected array, offering visual representations of intricate organization patterns (Figure 2C). These profiles can reveal distinct features, such as uniform head to tail organization of monomers within array (Example 1), to a slightly irregular (Example 2), and further to a highly discontinuous structure (Example 3). This approach elucidates the variability of array organization and the annotations are compatible with any genome browser such as Geneious and IGV (Figure 2D) to help explain and understand the different patterns of satDNA organization.

### 2. SatDNA Analysis

#### 2.1. Clustering analysis for uncovering satDNA monomer relationships

In the identification of satDNA variants, the standard phylogeny tree approach encounters limitations, particularly in distinguishing very subtle differences among satDNA monomers together with slow processing speed in drawing and analysing trees as well as often found low bootstrap values, long run times and complex dendrogram organizations (Suppl. Figure 3). To address these challenges, we employ a dimensionality reduction-based method of satDNA monomer analysis (Figure 3A). This method facilitates a thorough exploration of satDNA variation by processing all monomers from the same genome simultaneously with huge efficiency. Two main algorithms used are PCA (Figure 3A, upper panels) and UMAP (Figure 3A, bottom panels), operating on the distance matrices generated from the alignments. The output are graphs with satDNA monomers represented with individual dots that can be coloured based on different attributes, such as different arrays, chromosomes or species. The information contained in clusters, or lack thereof, can provide valuable insight into evolution and genome organization of specific satDNA family. These visualizations can reveal specific satDNA characteristics based on the clustering patterns or their absence. Certain satDNA monomers may show distinct segregation (Figure 3A, Example 1), indicating sequence variability and putative evolutionary events. Furthermore, it is possible to observe satDNA with a high degree of intrachromosomal similarity that nevertheless exhibits some degree of interchromosomal mixing (Figure 3A, Example 2) indicating several distinct expansion events. There are also instances where satDNA has undergone pronounced intrachromosomal exchanges (Figure 3A, Example 3), indicating active and dynamic genomic interactions within and between chromosomes. SatXplor provides interface for running either statistical (PCA) and geometric approaches (UMAP) for dimensionality reduction showing similar results, however the results vary between satDNA families thus it is best to run them both.

**Figure 3.**
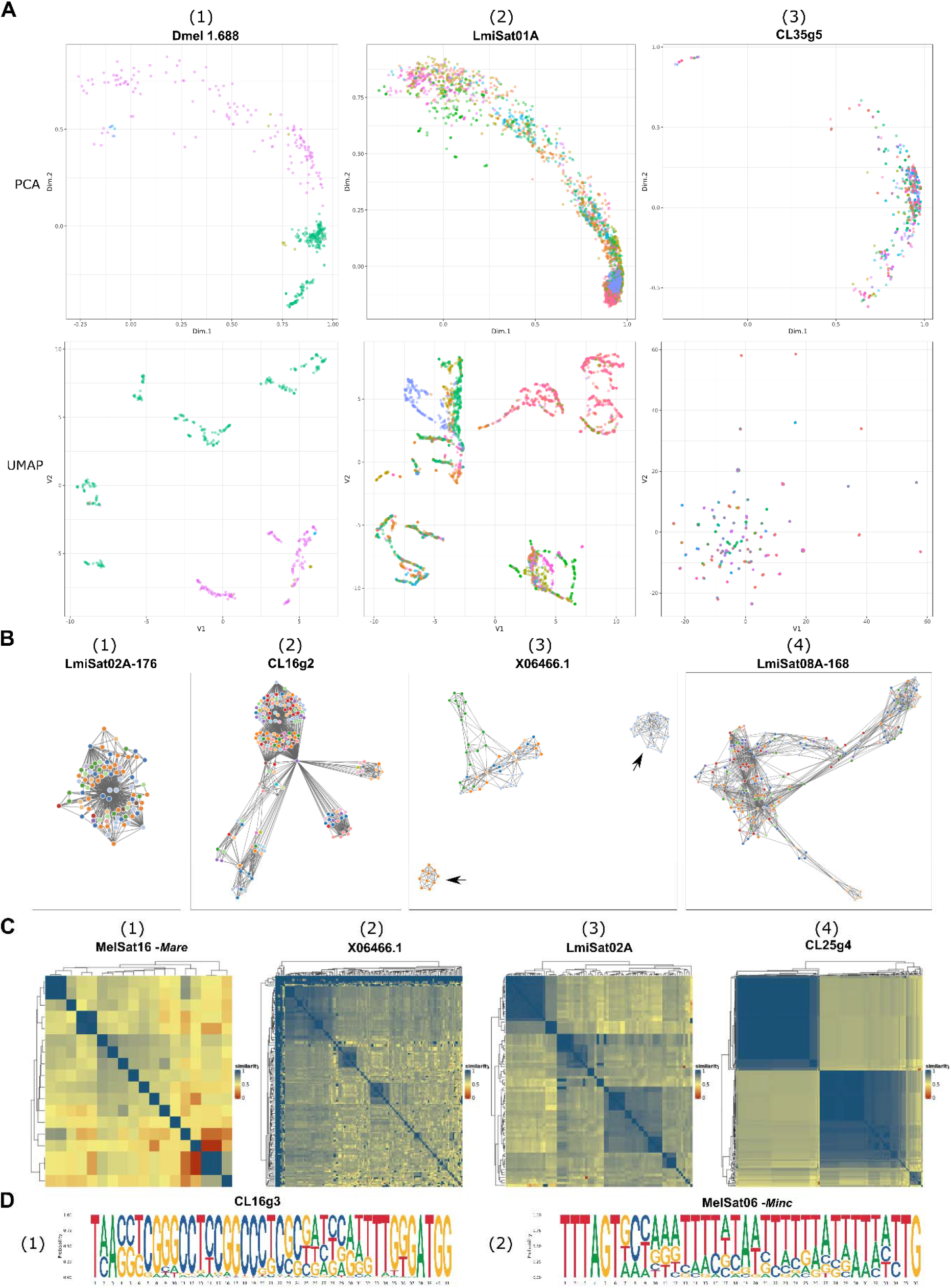
Evolution and organization analysis outputs of SatXplor. **A SatDNA monomer clustering analysis using PCA and UMAP.** The monomers are coloured according to a chromosome of origin. Specific satDNA monomers can show clear separation on both PCA (upper panel) and UMAP (bottom panel) (1), intrachromosomal similarity with certain degree of mixing with other chromosomes can sometimes be visualized only with one of the algorithms, in this case UMAP (2) and extremely high degree of intra- and intrachromosomal exchange that prevents efficient clustering by the algorithms (3). **B Graph networks**. Graph networks of arrays for four different satDNAs based on their sequence similarity relationship. Each dot on these graphs represents an array coloured based on the chromosome they originate from. SatDNA arrays with high degree of mixing represented by dense cluster of closely connected arrays (1), several arrays ‘ clusters that are distant from the central region (2), separated arrays ‘ clusters (marked with black arrows) (3) and example where there are many arrays ‘ clusters almost all linked together (4). **C SatDNA array surrounding region analysis**. Pairwise sequence similarity matrices of 500 bp neighbouring regions satDNA arrays visualized with *pheatmap*. Neighbouring regions of satDNA arrays without similarity (1), example of several small blocks indicating limited correlation with several sequences (i.e. transposons, other satDNAs). (2), mixture of different conserved sequences in the neighbouring regions represented by blocks of different sizes (3) and satDNA flanking regions that exhibit conserved large blocks indicating a conserved pattern of colocalization (4). **D Microhomology analysis**. Analysis of flanking regions can reveal the presence of high GC-content in a usually AT-high environment of satDNA (1) or regions that have poly(A/T) stretches. All satDNA names and species used (A-D) are listed in Suppl. Table 2.

#### 2.2. Unveiling evolutionary trends trough satDNA network analysis

Finally, SatXplor uses undirected graph networks to explain satDNA-specific evolutionary history. To achieve this, a specific algorithm was developed that creates an interactive, undirected graphs containing all annotated satDNA arrays based on genetic distance. This approach reveals the relationships and connections between specific arrays, whereas dimensionality reduction methods highlight the most likely scenario for the evolution and spread of a particular satDNA sequence in the genome. In the graph networks, various patterns emerge based on the distribution and clustering of satDNA monomers (Figure 3B). Arrays exhibiting a high degree of mixing are depicted by one cohesive cluster (Figure 3B, Example 1), Conversely, there are clusters distinct from the central region (Figure 3B, Example 2) or even separated (Figure 3B, Example 3), indicating divergence of arrays over time with mixed scenarios characterized by numerous interconnected clusters (Figure 3B, Example 4).

#### 2.3. Distance mapping of surrounding regions

SatDNA may be associated with other repetitive elements or embedded in gene-rich regions. Therefore, it is essential to assess satDNA features and recognize potential genomic association with different repeats or specific sequences. After the precise determination of the array edges (see Results section 1.4), the contiguous regions of 500 bp up- and downstream surrounding each array are systematically extracted. The size of investigated regions can be varied during explorative phase and later adjusted depending on the attributes of particular satDNA array. A detailed distance map is then generated based on the multiple alignment for each of these regions. Analysing the distance maps can reveal various patterns in satDNA surrounding regions (Figure 3C). One of them shows satDNAs with arrays that have almost no similarity between their neighbouring regions (Figure 3C, Example 1). The other pattern of satDNA arrays surrounding regions show high similarity regions at arrays edges representing occasional mixing with some other repeats (Figure 3C, Example 2). Interestingly, some satDNA arrays exhibit more frequent mixing with other highly similar or repetitive regions (Figure 3C, Example 3) indicating the presence of several distinct sequences in the vicinity of satDNA arrays. Finally, some arrays show large, conserved blocks (Figure 3C, Example 4) that originate from highly conserved sequences in the vicinity. In some cases, this algorithm can also detect the presence of sequences with shared homology only on one side of the array, suggesting possible association with other repeats and not a uniform embedding within repetitive regions.

#### 2.4. Microhomology detection in flanking regions of satDNA arrays

Additionally, precise edge detection algorithm allows SatXplor to focus on microhomologies near satDNA arrays. Microhomologies have been proposed to mediate both microhomology induced break repair and eccDNA genome reintegration (reviewed in Yang et al., 2022), and as such represent a vital pathway in evolution of satDNA in the genome (Sproul et al., 2020). The sequence logo graphs can be used to spot potential conserved microhomology regions (Figure 3D) as targets for investigation and manual curation or to reject certain mechanisms in organisms of interest. It allows finding specific satDNA characteristics, such as high GC content (Figure 3D, Example 1) within repetitive environments that are typically AT-rich. Additionally, microhomology analysis can detect regions containing polyN (A, T, G, C) stretches (Figure 3D, Example 2) indicative of potential regulatory role of sequence motifs.

## Discussion

The study of repetitive DNA poses a particular challenge, especially in the context of long stretches of satDNA, which have major structural and evolutionary implications (Louzada et al., 2020). There is a huge struggle to discern and accurately represent repetitive regions, leading to gaps and misrepresentations in genome assemblies and often requiring the implementation of multiple tailored approaches to obtain suitable platforms for their investigation (Peona et al., 2021). While achieving contiguous assemblies represents a significant milestone in satDNA research, the formidable challenge lies in the development of algorithms capable of accurately describing and deeply analysing these repetitive elements within the complexities of the genome.

SatXplor pipeline is a set of algorithms developed for precise definition and comprehensive analysis of satDNA in the assemblies of complex genomes. An assembly based on long reads and enriched in satDNAs is used as platform in analyses with SatXplor pipeline. This approach enables the precise characterization of satDNA monomer variability, which can range from very homogeneous to highly variable, providing specific parameters for their annotation. It facilitates the identification of how the satDNA monomers are arranged within arrays, distinguishing between typical tandems, complex organization of satDNA arrays and the presence of monomers in different orientations. In addition, SatXplor uses high-throughput analysis of all monomers of particular satDNAs with PCA and UMAP visualization to uncover underlying patterns in satDNA evolution. It is the first tool to define and precisely annotate both satDNA arrays and their edges enabling further assessment of the genomic environment, presence of conserved sequences, association with mobile elements or euchromatic regions. Moreover, SatXplor provides insights into the exact sites of satDNA insertion, leveraging microhomology evaluation to pinpoint insertion sites with precision. Graph networks show connections and distances among the arrays, offering valuable insights into evolutionary dynamics of satDNAs on the genome scale. SatXplor pipeline can also be used as a complementary pipeline following any algorithm utilized for the detection of repetitive elements.

SatXplor is optimized for satDNA with low to moderate genome abundancies (<5%) and monomer lengths 80-1000 bp. Running the pipeline for micro- and mini-satDNA may require manual parameter adjustments and verification. This pipeline proved particularly efficient in analyses of euchromatic satDNA, a distinct class of tandem repeats mainly localized in gene-rich regions of the genome which is characterized by moderate array lengths, variable copy numbers and a propensity for interspersion with protein-coding genes (Volarić et al., 2024). While the exact functions of euchromatic satDNAs are still unknown, their potential roles include involvement in genome stability, regulation of gene expression and chromatin organization, with specific evolutionary mechanisms influencing their propagation dynamics (Sproul et al., 2020). SatXplor emerges as a versatile tool for satDNA analysis, featuring compatibility with various detection tools and requiring only satDNA monomer consensus sequence and genome assembly sequences as input. A notable strength of SatXplor is the efficient detection of satDNA variation optimized for fast processing of all monomers within the genome. The pipeline streamlines the entire process by unifying the explorative and evolutionary analysis of satDNA in a single pipeline. It introduces a fast k-mer counting algorithm for array edge definition, which contributes to a vastly improved accuracy (max 5 bp, previously +/-monomer length) in satDNA array definition. Additionally, SatXplor allows exploration of neighbouring array regions, reveals both micro- and macrohomologies together with interactive graph networks providing insights into possible mechanisms of both intra- and interchromosomal spread and evolution.

Running SatXplor on different genome assemblies has provided valuable insights which can serve as a great starting point for in-depth analysis of satellitomes. It managed to successfully analyse even some centromeric and highly abundant satDNA, as demonstrated by identifying chromosome specificity of satDNA in D. melanogaster and *L. migratoria*. Additionally, it was able to detect conserved flanking regions around satDNA arrays in *M. incognita* and *A. thaliana*.

Although there are numerous tools for the detection of satDNA sequences, only a handful of algorithms are specifically designed for intragenomic satDNA analysis. However, most of these programs lack the versatility offered by SatXplor. For example, TRAP is an algorithm with both detection and analysis capabilities but limited to only parsing TRF outputs (Sobreira et al., 2006). SatDNA Analyzer recognizes intraspecies variation, detects polymorphisms and generates consensus sequences that provide a valuable starting point for phylogenetic studies (Navajas-Pérez et al., 2007), but requires multi species input in order to perform analysis. There are also programs like RepeatAnalyzer, focused only on intergenic microsatellite tracking, management, analysis and cataloguing (Catanese et al., 2016), however it is developed exclusively on bacterial data. There are also some programs with specific functions such as TRTTools with a focus on TR genotyping (Mousavi et al., 2021) or RepeatOBserver (Elphinstone et al., 2023) and StainedGlass (Vollger et al., 2022) with focus on visualization of centromeric repeats. As shown, customized programs are created usually only for specific targets, underscoring the need for a unified, versatile tool that encompasses both detection and downstream analysis. SatXplor bridges this gap by offering an integrated solution for satDNA exploration and analyses, providing a new approach in the field of repetitive DNA and setting a new standard for comprehensive satDNA studies. Importantly, its utility extends to functional satDNA analysis, enabling the identification of potential mechanisms and evolutionary trends, thereby advancing our understanding of genome dynamics.

## Materials and methods

### Genome and satDNA data

SatXplor algorithms were developed and tested on genomes of *T. castanuem, D. melanogaster, L. migratoria, M. incognita, M. arenaria* and *A. thaliana* with accessions provided in Suppl. Table 1. Information on analysed satDNA sequences as well as the sources are provided in Suppl. Table 2.

### SatXplor pipeline overview

#### Detection and extraction of monomers

The process of monomer detection based on query satDNAs is performed by NCBI BLAST+ (Camacho et al., 2009), running from a python subprocess command. The program also creates the subject database and deletes it since it is a fast process for most genomes. The main parameters for the BLAST search are:

> ‘-*evalue 10* –*outfmt 6* –*max*_t*arget*_*seqs 10000* –*task blastn –num_threads 2* -*dust no* -*soft_masking false* ‘

Dusk and soft masking are turned off in order to ensure detection of low complexity satDNA sequences.

#### Array creation

Arrays for a single satDNA family were created using a custom python script. First all BLAST hits from a single satDNA family were ordered by chromosome and their respective start position. Next, absolute distances were calculated between consecutive monomers. The resulting distances were then graphically visualized, and a histogram of the data was constructed. Since the histogram is a one-dimensional array, peaks were identified using *scipy*.*findpeaks()* function, keeping only peaks which amount to >5% of total monomer number in the genome (meaning that at least 5% of monomers are peak distance apart). Afterwards the max peak value for each family was used as an “extension factor “ by which each monomer annotation end position was elongated and finally each overlapping groups of monomer annotations were found and annotated as arrays.

#### Monomer density profiles

The 2D density approximations from BLAST output were created using a custom python script. First a 2D probability density function was done for both approximate query coverage and percentage identity using *scipy*.*stats*.*gaussian_kde()*. Then the data was binned in approximate bins by their total number determined by NKERNEL_BINS parameter and the values were plotted.

#### Kmer-based profiling of arrays

To find the exact edges of individual arrays, SatXplor employs the following algorithm for each satDNA under examination:

1. Extract all monomers form the genome
2. Create synthetic monomer-dimers
3. Extract extended arrays (arrays with +/-500bp flanks)
4. Create a hash table of all 32 k-mers from both the synthetic dimers and the extended arrays
5. For each 32 k-mer in the extended array, find the closest 32 k-mer from the synthetic dimer table by Hamming distance
6. Calculate a rolling sum score by averaging +/-5bp scores for each position
7. Define new array edges as the positions where first/last k-mer in the window has a score lower/higher than 5 (meaning an 87% similarity to a hit in the database).

#### Alignment and distance calculations

The main algorithm used in steps of monomer, flank and microhomology alignment is MAFFT (Katoh et al., 2002) which is run through a python subprocess command with the main settings:

> ‘ *mafft* --*adjustdirection* –*reorder* --*threads str(half*_*threads)* ‘

Where by default *half*_*threads* are half of total CPU processors available. Subsequent genetic distances are calculated by ape package in R using the “F81 “ genetic distance model.

#### Dimensionality reduction

The Principal Component Analysis (PCA) and Uniform Manifold Approximation and Projection (UMAP) algorithms and their implementation in R (R Core Team, 2022) were used to reduce the dimensionality of the distance matrices generated from the alignment steps. For the PCA implementation in R, the *PCA()* function from the *FactoMineR* (Lê et al., 2008) package was used. The resulting principal components are then visualized in reduced-dimensional space using the *ggplot2* (Wickham, 2016) package and the Scree plots for the first 10 principal components are also created. For the UMAP implementation in R, the *umap()* function from the umap package was used to find the UMAP embedding of the distance matrices with default parameters controlling aspects such as the number of dimensions in the reduced space and the number of neighbours to be considered for the local structure. The resulting UMAP embedding are also visualized directly with the *ggplot2* package.

#### Network and distance graphs

In our analysis pipeline, we used the *pheatmap* package in R to generate distance maps from the distance matrices of the flanking region alignments. To visualize networks, we used the following steps:

1. Calculate the distance matrix from the alignment of all monomers in the genome
2. Find the closest N monomers not belonging to the same array
3. Create a table off all possible array-array connections
4. Remove redundant or duplicate connections
5. Create a regular network using *igraph*
6. Create HTML interactive networks using *network3D*

### Performance

The performance of SatXplor is mainly defined by two key factors: number of input satDNA sequences and their respective number of monomers, with the greatest emphasis being on the total number of monomers/arrays present in the genome. All the examples were run on an Intel(R) Core(TM) i9-9900 CPU with 128GB RAM, but all the programs are configured to run on any machine >8GB of RAM and a multicore processor, however the run times may differ.

**Table 1.** Examples of run times, CPU and memory usage for the five organisms we used in testing the pipeline.

| Genome (size) | Total monomers | Peak memory (GB) | Wall time (seconds) |
| --- | --- | --- | --- |
| *Arabidopsis thaliana* (119 Mb) | 12107 | 0.6 | 230 |
| *Drosophila melanogaster* (143 Mb) | 426 | 0.5 | 120 |
| *Meloidogyne incognita* (199 Mb) | 10400 | 0.8 | 344 |
| *Meloidogyne arenaria* (297 Mb) | 83460 | 8 | 626 |
| *Tribolium castaneum* (225 Mb) | 11712 | 1 | 362 |
| *Locusta migratoria* (6.2 Gb) | 148132 | 25 | 5238 |

In general, it is advisable not to use the most abundant satDNA sequences in large genomes with this pipeline, as they may contain hundreds of thousands of monomers in the assembly and are likely to take a long time to complete. It is important to note that most of the pipeline is multithreaded at points where the likelihood of an I/O bottleneck is low. However, when using a large machine (e.g. a cluster) with a relatively slow drive, it is best to limit the number of threads used.

## Author Contributions

Conceptualization and Validation: M.V., N.M. and E.D.-S.; Formal Analysis, Investigation, Methodology, Visualization: M.V. and E.D.-S; Resources: N.M.; Software: M.V.; Supervision: E.D.-S. and N.M, Writing—Original Draft Preparation: E.D.-S; Writing—Review and Editing, M.V., N.M. and E.D.-S; Funding Acquisition: N.M. All authors have read and agreed to the published version of the manuscript.

