## Supplementary Material for "SatXplor – A comprehensive pipeline for satellite DNA analyses in complex genome assemblies"

**Supplementary Table 1.** List of used genomes, their sizes and NCBI GenBank accessions or source publications used in testing SatXplor

| Species | Total Length (Mbp) | Accession |
| --- | --- | --- |
| *Tribolium castaneum* | 225 | GCA_950066185 |
| *Arabidopsis thaliana* | 119 | GCF_000001735.4 |
| *Locusta migratoria* | 6200 | GCA_026315105.1 |
| *Meloidgyne incognita* | 199 | https://doi.org/10.57745/RAZ8JS |
| *Meloidgyne arenaria* | 297 | https://doi.org/10.57745/VBCSTD |
| *Drosophila melanogaster* | 142 | GCF_000001215.4 |

**Supplementary Table 2.** List of used satDNA names and NCBI GenBank accessions or source publications used in testing of SatXplor.

| Species | SatDNA Names | MonomerLength (bp) | Source |
| --- | --- | --- | --- |
| *Arabidopsis thaliana* | AR11 | 198 | X06466.1 (Simoens et al. 1988) |
| *Arabidopsis thaliana* | AR12 | 179 | X06467.1 (Simoens et al. 1988) |
| *Arabidopsis thaliana* | AR13 | 178 | X06468.1 (Simoens et al. 1988) |
| *Arabidopsis thaliana* | AR21 | 496 | X06470.1 (Simoens et al. 1988) |
| *Arabidopsis thaliana* | AR14 | 177 | X06469.1 (Simoens et al. 1988) |
| *Arabidopsis thaliana* | AR22 | 503 | X06471.1 (Simoens et al. 1988) |
| *Arabidopsis thaliana* | AR3 | 481 | X06472.1 (Simoens et al. 1988) |
| *Arabidopsis thaliana* | Clone 164A | 807 | X92080.1 (Simoens et al. 1988) |
| *Meloidogyne sp.* | Melsat1-83 | 48-295 | (Despot-Slade et al. 2022) |
| *Locusta migratoria* | LmiSat01A-185 | 370 | (Ruiz-Ruano et al. 2016) |
| *Locusta migratoria* | LmiSat02A-176 | 352 | (Ruiz-Ruano et al. 2016) |
| *Locusta migratoria* | LmiSat08A-168 | 336 | (Ruiz-Ruano et al. 2016) |
| *Locusta migratoria* | LmiSat14A-216 | 432 | (Ruiz-Ruano et al. 2016) |
| *Locusta migratoria* | LmiSat35A-228 | 456 | (Ruiz-Ruano et al. 2016) |
| *Locusta migratoria* | LmiSat54A-272 | 544 | (Ruiz-Ruano et al. 2016) |
| *Tribolium castaneum* | Cast1-9 | 121-350 | (Pavlek et al. 2015; Volarić et al. 2024) |
| *Drosophila melanogaster* | Dmel1688 | 350 | KY575282.1 (Khost, Eickbush, and Larracuente 2017) |
| *Meloidogyne incognita* | CL16g1 | 83 | (Despot-Slade et al. 2021) |
| *Meloidogyne incognita* | CL16g2 | 50 | (Despot-Slade et al. 2021) |
| *Meloidogyne incognita* | CL16g3 | 45 | (Despot-Slade et al. 2021) |
| *Meloidogyne incognita* | CL25g4 | 77 | (Despot-Slade et al. 2021) |
| *Meloidogyne incognita* | CL32g5 | 44 | (Despot-Slade et al. 2021) |

**Supplementary Table 3.**  BLAST output table example. The shaded cells (perc_id and al_len) are used for 2D kernel density approximations. “s_start” and “s_end” columns are used for retrieving exact annotations in the genome.

| query | subject | perc_id | al_len | MM | GO | s_start | s_end | evalue | score |
| --- | --- | --- | --- | --- | --- | --- | --- | --- | --- |
| MelSat01-295 | Marev4contig205 | 96.61 | 295 | 10 | 0 | 50823 | 50529 | 8.69E-137 | 488 |
| MelSat01-295 | Marev4contig205 | 96.61 | 295 | 10 | 0 | 108058 | 107764 | 8.69E-137 | 488 |
| MelSat01-295 | Marev4contig205 | 96.61 | 295 | 8 | 1 | 3519 | 3227 | 1.06E-135 | 485 |
| MelSat01-295 | Marev4contig205 | 96.61 | 295 | 8 | 1 | 4693 | 4401 | 1.06E-135 | 485 |
| MelSat01-295 | Marev4contig205 | 96.61 | 295 | 8 | 1 | 5652 | 5360 | 1.06E-135 | 485 |
| MelSat01-295 | Marev4contig205 | 96.61 | 295 | 8 | 1 | 8891 | 8599 | 1.06E-135 | 485 |
| MelSat01-295 | Marev4contig205 | 96.61 | 295 | 8 | 1 | 14470 | 14178 | 1.06E-135 | 485 |
| MelSat01-295 | Marev4contig205 | 96.61 | 295 | 8 | 1 | 15938 | 15646 | 1.06E-135 | 485 |
| MelSat01-295 | Marev4contig205 | 96.61 | 295 | 8 | 1 | 16231 | 15939 | 1.06E-135 | 485 |
| MelSat01-295 | Marev4contig205 | 96.61 | 295 | 8 | 1 | 17990 | 17698 | 1.06E-135 | 485 |

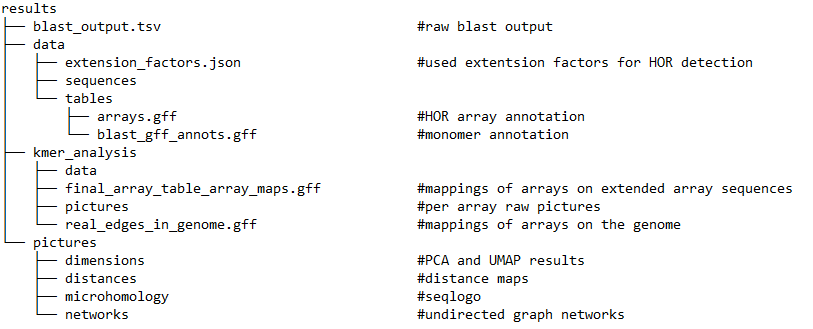
**Supplementary Figure 1.** Structure of the SatXplor result output directory.

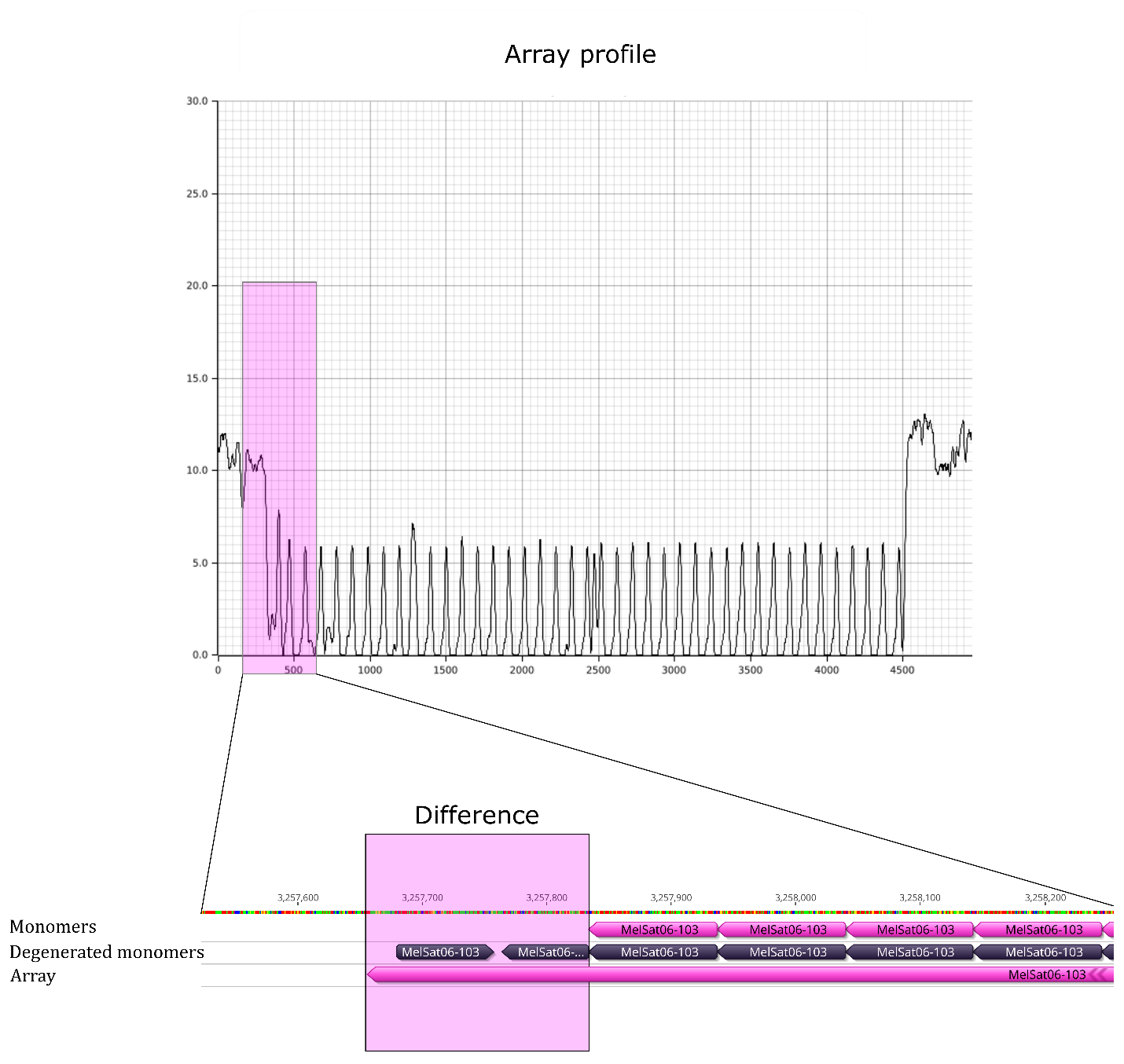

**Supplementary Figure 2.** Schematic of the effect of k-mer based edge detection on proper identification of array edges. This is a representative example for MelSat06-103 satDNA in *M. arenaria*. The upper part of the figure depicts the k-mer distance profiles, while the lower shows the accompanying schematics of annotation improvement. In genome tracks, the monomers annotated by BLAST are in the first and second row (Monomers and Degenerated monomers) while the newly annotated array-edges are in the third row (Arrays). If only the Monomers track is used for the flanking satDNA analysis, the surrounding array region would still correspond to Degenerated monomers, making it impossible to find the exact region of satDNA insertion. Using the kmer-based approach solves this problem, and the new edge is better detected, with a difference of almost 100 bp.

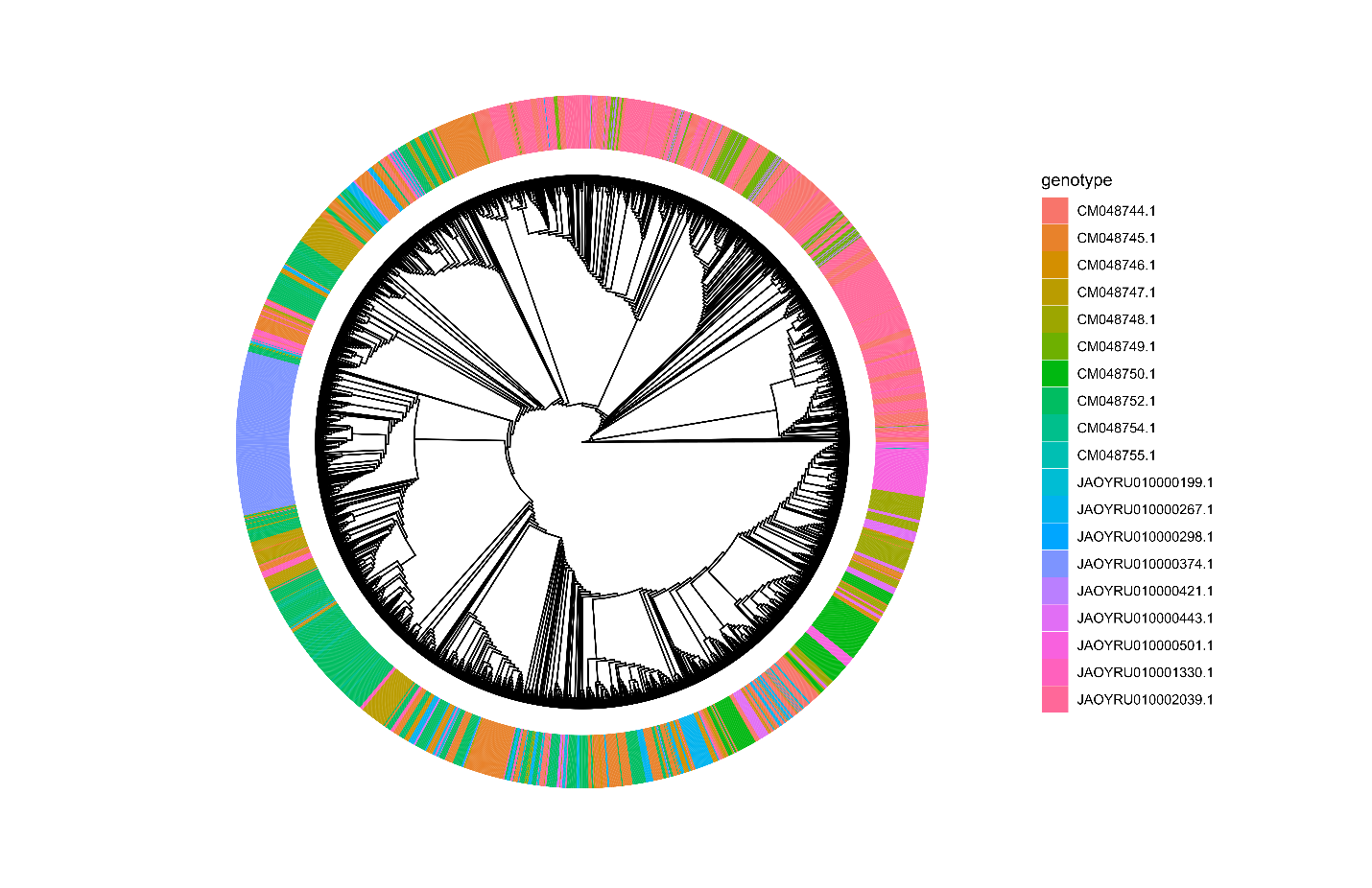
**Supplementary Figure 3.** IQ-TREE generated tree of the *L. migratoria* all extracted LmiSat01A-185 monomers (3512). Colors are based on the chromosome of origin with hard to visualize precise differences and extremely large tree. Overall groupings are similar to the UMAP results (Figure 3A, Example 2), however the runtime was 8x longer than the whole SatXplor pipeline run for all satDNA sequences in the genome (14h2m vs 1h26m). IQ-TREE was run with the MAFFT generated alignment and *–B 1000* settings.
